# A cell-free framework for biological systems engineering

**DOI:** 10.1101/018317

**Authors:** Henrike Niederholtmeyer, Zachary Z. Sun, Yutaka Hori, Enoch Yeung, Amanda Verpoorte, Richard M. Murray, Sebastian J. Maerkl

## Abstract

While complex dynamic biological networks control gene expression and metabolism in all living organisms, engineering comparable synthetic networks remains challenging^1,2^. Conducting extensive, quantitative and rapid characterization during the design and implementation process of synthetic networks is currently severely limited due to cumbersome molecular cloning and the difficulties associated with measuring parts, components and systems in cellular hosts. Engineering gene networks in a cell-free environment promises to be an efficient and effective approach to rapidly develop novel biological systems and understand their operating regimes^3-5^. However, it remains questionable whether complex synthetic networks behave similarly in cells and a cell-free environment, which is critical for *in vitro* approaches to be of significance to biological engineering. Here we show that synthetic dynamic networks can be readily implemented, characterized, and engineered in a cell-free framework and consequently transferred to cellular hosts. We implemented and characterized the “repressilator”^6^, a three-node negative feedback oscillator *in vitro*. We then used our cell-free framework to engineer novel three-node, four-node, and five-node negative feedback architectures going from the characterization of circuit components to the rapid analysis of complete networks. We validated our cell-free approach by transferring these novel three-node and five-node oscillators to *Escherichia coli*, resulting in robust and synchronized oscillations reflecting the *in vitro* observation. We demonstrate that comprehensive circuit engineering can be performed in a cell-free system and that the *in vitro* results have direct applicability *in vivo*. Cell-free synthetic biology thus has the potential to drastically speed up design-build-test cycles in biological engineering and enable the quantitative characterization of synthetic and natural networks.

## MAIN

A central tenet of engineering involves characterizing and verifying prototypes by conducting rapid design-build-test cycles in a simplified environment. Electronic circuits are tested on a breadboard to verify circuit design and aircraft prototypes are tested in a wind tunnel to characterize their aerodynamics. A simplified environment does not exist for engineering biological systems, nor is accurate software based design possible, requiring design-build-test cycles to be conducted *in vivo*^2^. To fill the gap between theoretical design and laborious *in vivo* implementation for biological systems we devised a cell-free framework consisting of *E. coli* lysate (“TX-TL”)^5,7^ and a microfluidic device capable of emulating cellular growth and division^3^ (Fig. 1). Almost all prototyping can be done on linear DNA, which requires less than 8 hours to assemble and test. The cell-free framework provides a simplified and controlled environment that allows us to drastically reduce the design-build-test cycle^8^.

**Figure 1.**
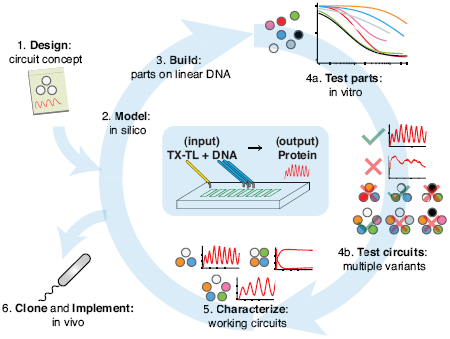
The cell-free framework allows rapid and extensive characterization of biological systems. Schematic representation of the design-build-test cycle using the cell-free framework. A design is first modeled to obtain intuition about the architecture. Parts are then assembled on linear DNA without cloning, and tested *in vitro*. With functional parts, circuit variants can then be tested and working circuits can be extensively characterized. Final circuits are cloned onto plasmids and implemented *in vivo*. Center shows the microfluidics device used. Input is a circuit encoded by linear or plasmid DNA and TX-TL *in vitro* reagent, which is then translated and transcribed into protein. For a specific example of the cell-free framework applied to engineering a 5-node oscillator network see Supp. Fig. 3.

We first asked whether our cell-free framework could be used to run and characterize an existing synthetic *in vivo* circuit and chose to test the repressilator^6^ as a model circuit. We successfully implemented the original repressilator network in our cell-free framework and observed long-term sustained oscillations with periods matching the *in vivo* study (Fig 2, Supp. Video 1). We compared the original repressilator to a modified version containing a point mutation in one of the CI repressor binding sites in the promoter regulating LacI (Fig. 2a). This mutation increases the repressor concentration necessary for half-maximal repression (K_M_), and reduces cooperativity^9^. At long dilution times (t_d_) both circuits oscillated, but with shifted absolute reporter protein concentrations (Fig. 2b). At decreasing dilution times amplitudes decreased and periods became faster with a linear dependence on t_d_. Faster dilution times, however, did not support oscillations for the modified network (Fig. 2b-c). Experimentally, the range of dilution times supporting oscillations can serve as a measure for robust oscillator function, which generally diminishes with decreasing synthesis rates or when binding of one repressor to its promoter is weakened as in the O_R_2* mutant (Supp. Fig. 1). To give another example for an experimental characterization that would be challenging to perform in a cellular environment we analyzed the repressilator network in phase space showing limit cycle oscillations and invariance to initial conditions (Fig 2d).

**Figure 2.**
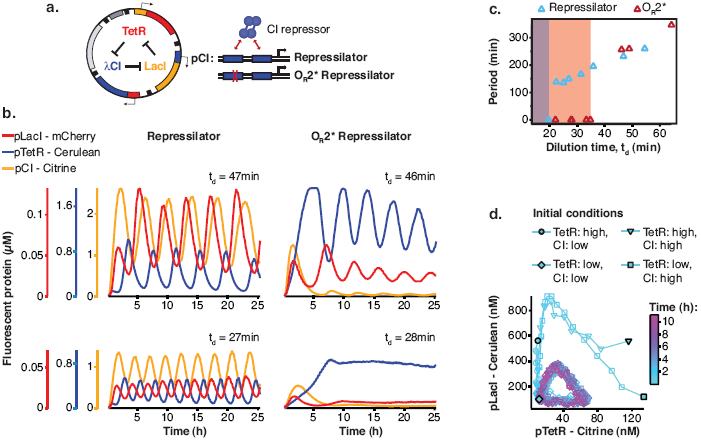
Cell-free repressilator characterization. **a**, Application of the cell-free framework to characterize the original repressilator^6^ and a modified version with a point mutation in the CI promoter (O_R_2*) located in one of the binding sites of the CI repressor. **b**, Expression from the three promoters of the repressilator and the O_R_2* version at different dilution times. **c**, Oscillation periods of the repressilator as a function of dilution time. In the O_R_2* version sustained oscillations were supported in a narrower range of dilution times as compared to the original repressilator network. **d**, Phase portrait of repressilator oscillations starting from different initial TetR and CI repressor concentrations.

The cell-free framework also allows rapid characterization of individual network components. We measured the transfer functions of repressor-promoter pairs in the repressilator network (Fig. 3a, Supp. Fig. 2a,b, Table 1) and found that the network is symmetric in terms of transfer functions. In the CI promoter O_R_2* mutant we observed the expected shift in K_M_ and decreased steepness of the transfer function. We also characterized TetR repressor homologs as building blocks for novel negative feedback circuits (Fig 3a) and with the exception of QacR observed similar transfer functions as observed *in vivo*^10^ (Supp. Fig 2c).

**Figure 3.**
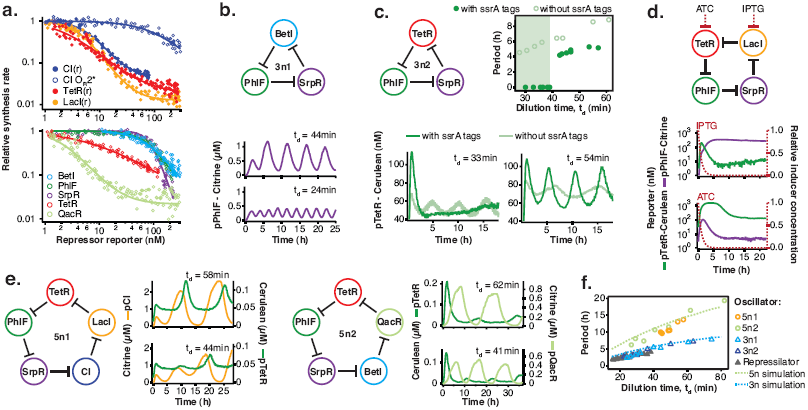
Cell-free prototyping and characterization of novel negative feedback circuits. **a**, Transfer functions of the repressilator repressor-promoter pairs (top) and TetR homologs (bottom). The TetR repressor was tested against two different promoters: the promoter used in the repressilator (top panel) and the J23119-TetR promoter^10^ (bottom panel). Lines are Hill function fits. **b**, Oscillations of a novel 3-node ring oscillator (3n1) constructed on plasmid DNA. **c**, Two versions of a second 3-node ring oscillator (3n2) on linear DNA were used to study the effect of ClpXP degradation on oscillator function. One version was ssrA-tagged on all repressor genes while the other version did not carry degradation tags on the repressors. The same reporter with a medium-strength degradation tag was used in both versions. **d**, A 4-node cyclic negative feedback network on linear DNA has two stable steady states that depend on the initial conditions. IPTG switched the network into the state where pPhlF was on and pTetR off. An initial pulse of aTc resulted in the opposite stable steady state. **e**, Two 5-node negative feedback architectures oscillated with longer periods than our 3-node networks as predicted by simulations (Supplementary Information) (f).

Using three new repressors, BetI, PhlF and SrpR, we constructed a novel 3-node (3n) circuit (3n1) and observed high-amplitude oscillations over a broad range of dilution times with the same dependence of amplitude and period on t_d_ as for the repressilator (Fig 3b). In our characterization of the repressilator network and the 3n1 oscillator we found dilution rates to be critical for the existence, period and amplitude of oscillations. Protein degradation is similar to dilution in that it results in removal of repressor proteins. In order to study the effect of degradation we constructed a second 3n network (3n2) using TetR, PhlF and SrpR repressors on linear DNA. One version of the circuit used strong ssrA ClpXP degradation tags, while the second used untagged repressors. We observed oscillations for both circuits (Fig 3c). However, the circuit without ssrA-tag mediated protein degradation exhibited slower oscillations, which extended to lower dilution times, showing that protein degradation, just like dilution, affects oscillator function and period. Effects of ClpXP-mediated protein degradation, which have been shown to be important for existence and frequency of oscillations *in vivo*^11,12^, can thus be emulated in a cell-free environment.

Theory predicts that ring architectures built from an odd number of repressors oscillate, while even-numbered architectures have stable steady states^13,14^. We experimentally built and tested a 4-node circuit from LacI, TetR, PhlF and SrpR on linear DNA. Initial pulses of LacI inducer IPTG or TetR inducer aTc allowed us to switch expression into either one of the two stable steady states (Fig. 3d).

Encouraged by the robust oscillations observed in the 3n networks, we built two 5-node ring oscillators (5n) to test our prototyping environment on a novel synthetic network architecture (Fig. 3e). Despite their considerable complexity both circuits oscillated over a broad range of dilution times with the expected period lengthening, which could be as long as 19h. Comparing all ssrA-tagged 3n and 5n ring architectures, we show that the observed periods could be accurately predicted for all four networks by computational simulations (Fig. 3f). Our cell-free framework allows testing and characterization of complex networks including verifying networks cloned onto a single plasmid, which is the closest approximation to *in vivo* implementation (Supp. Fig. 3).

To validate our cell-free approach *in vivo* we cloned the 3n1 and 3n2 networks onto low-copy plasmids and co-transformed each with a medium-copy reporter plasmid into *lacI*-JS006 *E. coli*^15^. When tested on a microfluidic device (mother machine^16^), both 3n oscillators showed regular oscillations with periods of 6 ±1 hours for at least 30 hours (Fig. 4a, Supp. Video 2,3). Both oscillators were surprisingly robust as all cells undergoing healthy cellular division oscillated (n = 71) (Supp.. Fig. 4, **Supp. Video 4**).

**Figure 4.**
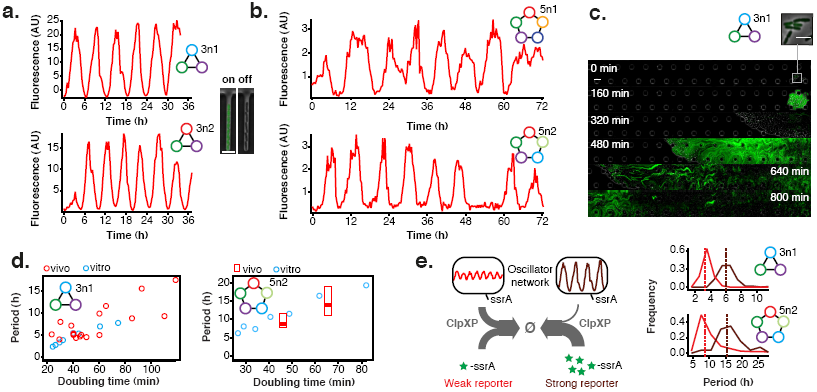
Novel 3-node and 5-node ring oscillators *in vivo*. **a**, Time series traces of 3-node ring oscillators running in *E. coli* (mother machine). Single trap traces of 3n1 and 3n2 observed for 36 h *in vivo* using a strong pPhlF sfGFP-ssrA reporter and a representative image from an “on” and “off” state of oscillation. Scale bar: 5 μm. **b**, Time series traces of 5-node ring oscillators running in *E. coli* (mother machine). Single trap traces of 5n1 and 5n2 observed for 72 h *in vivo* using a weak pPhlF sfGFP-ssrA reporter. **c**, 3n1 displays phase synchrony *in vivo* (CellASIC). Time series micrographs of 3n1 under a strong pPhlF sfGFP-ssrA reporter every 160 min; inset shows individual cells of the initial microcolony. Scale bar: 10 μm and 5 μm (inset). **d**, Relationship between period and division time *in vivo*. Left, 3n1 *in vivo* under a strong pPhlF sfGFP-ssrA reporter. The *in vitro* data is shown for comparison. Each point in the *in vivo* data corresponds to the period and division time from a CellASIC experiment run under different media type and flow rates. Right, 5n2 *in vivo* under a weak pPhlF sfGFP-ssrA reporter. *In vivo* periods determined at 29°C and 21°C growth temperature in mother machine experiments. Boxes represent the inner quartile range with the median. e, Influence of reporter concentration on oscillation periods by competing for ClpXP degradation. Left, with constant amounts of ClpXP the reporter concentration affects repressor degradation and thus oscillation period. Histograms of the periods observed with a weak and a strong pPhlF sfGFP-ssrA reporter for both 3n1 and 5n2 run in the mother machine. Dashed lines indicate the medians.

We next tested our 5n oscillators *in vivo*. Due to loading effects^17^, 5n1 was not viable when co-transformed with a strong reporter. When tested with a low expression strength reporter both 5n oscillators showed robust oscillations that were maintained for at least 70 hours, and over 95% of all analyzed traps containing healthy cells oscillated (n = 104). In addition, both 5n networks oscillated with similar periods: 8 hours for 5n1, and 9 hours for 5n2 (Fig. 4b, Supp. Fig. 4, **Supp. Video 5, 6, 7**).

We also tested both 3n oscillators on a CellASIC system, which allows planar single-layer colony formation. In this system we observed a striking population level synchronization of daughter cells inheriting the oscillator state from their mother cells (Fig. 4c, Supp. Fig. 5a, Supp. Video 8,9). Synchronization was also apparent when using three different fluorescent reporters simultaneously (Supp. Fig. 5b, **Supp. Video 10**). We did not observe population level synchronization in the original repressilator, the O_R_2* mutant (Supp. Fig. 5c) nor the 5n networks. Synchronized oscillations were not reported with the original repressilator^6^, and have only been observed in oscillators using intercellular communication^18,19^. We hypothesize that the 3n1 and 3n2 synchronization is due to increased repressor concentrations as compared to the original repressilator network (Supp. Fig. 5d), which increases the inheritance of the period phenotype and minimizes the rapid de-synchronization expected from stochastic cellular protein fluctuations^20^. However, a quantitative characterization of the synchronization phenotype requires more in depth understanding of stochastic effects *in vivo*.

Because cells were synchronized, we were able to analyze the population as a whole to make general conclusions of oscillator behavior. We varied dilution time by using different media conditions and media flow rates, and found a direct relationship between division times and period, consistent with the *in vitro* data collected. Oscillation periods of the 5n oscillators were also consistent with our *in vitro* results and showed a similar dependence on doubling time (Fig. 4d).

Finally, we compared 3n1 and 5n2 with weak and strong reporters *in vivo* to analyze the effect of protein degradation on the oscillator period. We theorized that given a constant concentration of ClpXP, stronger reporters would result in more ClpXP loading, thereby slowing the period of oscillation. ClpXP is thought to influence oscillation dynamics *in vivo* in this manner^11^. We found that in the mother machine, both the period distributions of 3n1 and 5n2 showed this characteristic (Fig. 4e), which reflects our *in vitro* findings of differential –ssrA tag dependent period length (Fig. 3c).

We demonstrated the utility of our cell-free framework for biological systems engineering and component characterization. We observed some differences between the *in vitro* and cellular environment, particularly in the difficulty of predicting cellular toxicity and loading effects of the 5n oscillators *in vivo*. While more work is necessary describing and explaining differences between *in vitro* and *in vivo* environments^8,21^, the observed behavior of complex networks in our cell-free environment reflected network behavior *in vivo* well. The cell-free framework is thus a powerful emulator of the cellular environment allowing precise control over experimental conditions and enabling studies that are difficult or time consuming to perform in cells. With further developments in cell-free lysate systems and supporting technologies, the *in vitro* approach is posed to play an increasing role in biological systems engineering and provides a unique opportunity to design, build, and analyze biological systems.

## METHODS

### TX-TL reactions

Preparation of TX-TL was conducted as described previously^7^, but using strain “JS006” co-transformed with Rosetta2 plasmid and performing a 1:2:1 extract:DNA:buffer ratio. This resulted in extract “eZS4” with: 8.7 mg/mL protein, 10.5 mM Mg-glutamate, 100 mM K-glutamate, 0.25 mM DTT, 0.75 mM each amino acid except leucine, 0.63 mM leucine, 50 mM HEPES, 1.5 mM ATP and GTP, 0.9 mM CTP and UTP, 0.2 mg/mL tRNA, 0.26 mM CoA, 0.33 mM NAD, 0.75 mM cAMP, 0.068 mM folinic acid, 1 mM spermidine, 30 mM 3-PGA, 2% PEG-8000. For experiments utilizing linear DNA GamS was added to a final concentration of 3.5 μM^8^.

### Steady-state reactions

Experiments were performed in a microfluidic nano-reactor device as described previousely^3,7^ with some modifications to optimize the conditions for the lysate-based TX-TL mix. Reaction temperature was 33°C. Lysate was diluted to 2x of the final concentration in 5 mM HEPES 5 mM NaCl buffer (pH 7.2). The reaction buffer mix was combined with template DNA and brought to a final concentration of 2x. For a 24 h experiment 30 μl of these stocks were prepared. During the experiment, lysate and buffer/DNA solutions were kept in separate tubing feeding onto the chip, cooled to approximately 6°C, and combined on-chip. We ran experiments with dilution rates (μ) between approximately 2.8 and 0.5 h^−1^, which corresponds to dilution times, t_d_ = ln(2) μ^−1^, between 15 and 85 min. These were achieved with dilution steps exchanging between 7 and 25% of the reactor volume with time intervals of 7 to 10 min, which alternately added fresh lysate stock or fresh buffer/DNA solution into the reactors. Dilution rates were calibrated before each experiment. Initial conditions for the limit cycle analysis of the repressilator network were set by adding pre-synthesized repressor protein at the beginning of each experiment. For this, CI repressor (together with Citrine reporter) and TetR repressor (together with Cerulean reporter) were expressed for 2.5h in batch. On chip the initial reaction was mixed to be composed of 25% pre-synthesis reaction and 75% fresh TX-TL mix and repressilator template DNA. Then, the experiment was performed at a t_d_ of 19.2 ± 0.3 min. Initial conditions for the 4-node experiment were 2.5μM aTc or 250μM IPTG, and the experiment was performed at a t_d_ of 44.5 ± 0.9 min. DNA template concentrations used in steady-state reactions are listed in Supp. Table 4. Arbitrary fluorescence values were converted to absolute concentrations from a calibration using purified Citrine, Cerulean, and mCherry, which were prepared using previously published protocols utilizing a His6 purification method followed by size-exclusion chromatography and a Bradford assay to determine protein concentration^8^.

### DNA and strain construction

DNA was constructed using either Golden Gate Assembly or Isothermal Assembly. The original repressilator plasmid, pZS1^6^ was used as a template for initial characterization and for construction of the O_R_2* mutant. Transfer function plasmids were constructed by Transcriptic, Inc. For other plasmids, partial sequences were either obtained from Addgene^10^ or synthesized on gBlocks or ssDNA annealed oligonucleotides (Integrated DNA Technologies). For linear DNA, all DNA was constructed using previously published Rapid Assembly protocols on a “v1-1” vector^8^. Specific plasmids required secondary-structure free segments, which were designed by R2oDNA^23^. JS006^15^ was co-transformed with origin-of-replication compatible plasmids to create engineered strains. Specifically, negative-feedback oscillator units were cloned onto pSC101* low copy plasmids (ampR or kanR), while reporters were cloned onto colE1 medium copy plasmids (kanR or cmR). To modulate the reporter copy number, all experiments were conducted below 37°C^24^. Strain passage was minimized to avoid plasmid deletions due to the *recA+* nature of JS006 and the high complexity of oscillator plasmids or triple-reporter plasmid. Based on the *in vitro* and *in silico* results, we used strong transcriptional and translational^25^ units to maximize gain.

### Transfer function measurement

Transfer functions of the repressor – promoter pairs were determined in the nano-reactor device at a minimum of two different dilution times (Supp. Fig. 2). All tested promoters were cloned into a plasmid in front of a BCD7 ribosomal binding site and the Citrine open reading frame. A non-saturating concentration of 1nM plasmid was used in the experiment. The repressors were expressed from linear templates carrying the J23151 promoter and the BCD7 ribosomal binding site with time-varying concentrations, which were increased from 0 to 2.5nM and decreased back to 0 during the course of the experiment^3^. Simultaneously we expressed Cerulean as a reporter for the repressor concentration from a linear template at an identical concentration as the repressor template. From the concentration of the Citrine reporter we calculated the synthesis rate of the fluorescent protein over time using a model of steady state protein synthesis in the nano-reactor device^3^,

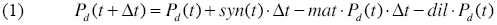

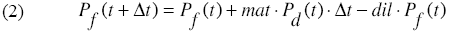

where *P*_*d*_ and *P*_*f*_ are dark and fluorescent reporter concentration respectively, t is time, Δt is the time interval between dilution steps, *dil* is the volume fraction replaced per dilution step, which was determined during the calibration of the device, and *mat* is maturation rate of the fluorescent protein. Maturation times of Citrine and Cerulean were determined as described previously^3^ and were 15 ±4 min for Cerulean and 29 ±3 min for Citrine. Dark fluorescent protein was calculated from equation (2):

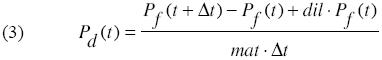

and the synthesis rate was calculated from equation (1):

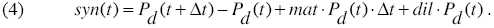

We used the sum of measured fluorescent Cerulean concentration and equation (3) for dark Cerulean as a measure of the total repressor protein present at any time during the experiment.

The synthesis rates were normalized to their respective maximal values (v_max_) and plotted against the concentration of the repressor reporter using only repressor concentrations higher than 1nM. The transfer curves were then fit to a Hill function

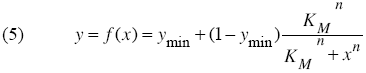

where y is the synthesis rate, *y*_min_ is the minimum synthesis rate, *n* is the Hill coefficient and *K*_M_ is the Michaelis Menten constant for half maximal promoter activity. The fitting was performed in Igor Pro using orthogonal distance regression with ODRPACK95 assuming a 9% error in the measurements of Citrine and Cerulean fluorescence.

### V_max_ measurements

Relative promoter strengths (v_max_ values) were determined using the transfer function promoter plasmids. *In vitro* strengths were determined in 5 μl TX-TL reactions at a DNA template concentration of 1nM. Reactions were assembled in 384-well plates, overlaid with 35 μl Chill-Out Liquid wax (BioRad) and analyzed using a Biotek SynergyMx plate reader set to 33°C reaction temperature, and reading Citrine fluorescence with Exc:510±9nm and Em:540±9nm. For comparison, Citrine fluorescence at 6h was normalized to the value of pLacI.

*In vivo* strengths were determined using *E. coli* JS006 transformed with the same plasmids. Cells were grown at 29°C in MOPS medium supplemented with 0.4% glycerol and 0.2% casaminoacids. For each strain, three independent overnight cultures were diluted 1:50 and grown to mid-log phase. They were then diluted to a starting OD_600_ of 0.15 into 100 μl growth medium in a 96-well plate and grown in the plate reader at 29°C with periodic shaking measuring Citrine fluorescence. Fluorescence values were normalized to OD resulting in steady state values after 2 h. Average steady state values were normalized to pLacI for comparison with the *in vitro* measurement.

### *In vivo* experiments

Mother machine^16^ experiments were conducted with custom-made microfluidic chips (mold courtesy of M. Delincé and J. McKinney, EPFL). *E. coli* cells were trapped in channels of 30 μm length, 2 μm width and 1.2 μm height. Before loading onto the device, cells were grown from a frozen stock to stationery phase. Cells were then concentrated 10-fold and loaded onto the chip. Experiments were performed using LB medium supplemented with 0.075% Tween-20 at a flow rate of 400 μl/h. Oscillation traces were collected from single mother machine traps using the background subtracted average fluorescence intensity of the entire trap.

CellASIC experiments were conducted using B04A plates (Merck Millipore, Darmstadt Germany). Flow rates were varied between 0.25 psi − 2 psi. Cells were grown from frozen stock in media at running temperature to stationery phase. Cells were then diluted 1:100 for 2 hours, and loaded on a equilibrated plate at 1:1000 or less to achieve single-cell loading efficiencies per chamber. To vary cellular doubling times, different growth media were used: LB (BD Biosciences), M9CA (Sigma Aldrich) with 0.2% glucose, 2xYT (MP Bio), MOPS EZ Rich (Teknova).

Cells were imaged in time series every 10-20 min using a 100x phase objective minimizing both lamp intensity (12% Xcite 120, Excelitas Inc. Waltam MA or 1-2% CoolLED pE-2, Custom interconnected Ltd., UK) and exposure times (<500ms) to limit photo-toxicity.

### Analysis of *in vivo* data

Images were processed and stitched^26^, if necessary, using Fiji/ImageJ^27^. Fluorescence traces of cell populations with synchronized oscillations were extracted from CellASIC movies using background corrected mean fluorescence intensity from the entire field of view. For cells that were not synchronized over the complete field of view, we tracked regions of oscillating sister cells at the edge of the microcolony. We used ImageJ to define polygonal regions around those cells and manually shifted the polygonal region to track the front of growing cells. Periods were determined from fluorescence traces derived from mother machine and CellASIC movies by measuring the time from one oscillation peak to the next peak. Doubling times were estimated by averaging over the doubling times of at least ten individual cells.

## Supplementary Information

This manuscript contains Supplementary Information: Supplemental Figures 1 – 5, Supplemental Videos 1 – 10, Supplemental Tables 1 – 5, Supplemental Model Information.

## Acknowledgements

We thank Yin He, Transcriptic, Inc. and Holly Rees for cloning assistance, Jan Kostecki and Stephen Mayo for protein purification and size exclusion chromatography assistance, Rohit Sharma and Marcella Gomez for initial testing and modeling of oscillators *in vitro*, Kyle Martin for laboratory assistance, Adam Abate, Tanja Kortemme, and Charles Craik for laboratory space and equipment, Matthieu Delincé, Joachim De Jonghe, Marc Spaltenstein, John McKinney and Jin Park for mother machine material and assistance, Tim Chang and Benjamin Alderete for CellASIC assistance, and Michael Elowitz for insights and scientific support. This work was supported in part by EPFL and the Defense Advanced Research Projects Agency (DARPA/MTO) Living Foundries program, contract number HR0011-12-C-0065 (DARPA/CMO). Z.Z.S. is also supported by a UCLA/Caltech Medical Scientist Training Program fellowship, Z.Z.S and E.Y by a National Defense Science and Engineering Graduate fellowship, and Y.H. by a JSPS Fellowship for Research Abroad. The views and conclusions contained in this document are those of the authors and should not be interpreted as representing officially policies, either expressly or implied, of the Defense Advanced Research Projects Agency or the U.S. Government.

## Author Contributions

All authors contributed extensively to the work presented in this paper.

## Author Information

The authors declare no competing financial interests. Correspondence and requests for materials should be addressed to S.J.M.

**Supplemental Table 1.** Transfer function parameters. Parameter values of repressor – promoter pairs were determined by fitting to the Hill equation as described in the Methods.

| **Name** | **K<sub>M</sub>** | **n** | **y<sub>min</sub>** |
| --- | --- | --- | --- |
| CI – pCI<sup>2</sup> | 5.9 | 1.9 | 0.04 |
| CI – pCI(O<sub>R</sub>2*)<sup>4</sup> | 103 | 1.2 | 0 |
| LacI – pLacI(r)<sup>2</sup> | 4.1 | 1.9 | 0.01 |
| TetR – pTetR(r)<sup>2</sup> | 2.3 | 1.2 | 0.02 |
| LacI – pLacI<sup>3</sup> | 3.6 | 1.7 | 0.01 |
| SrpR – pSrpR<sup>1</sup> | 86 | 3.6 | 0 |
| PhlF – pPhlF<sup>1</sup> | 79 | 2 | 0 |
| TetR – pTetR<sup>1</sup> | 7.6 | 0.8 | 0 |
| BetI – pBetI<sup>1</sup> | 75.4 | 1.7 | 0 |
| QacR – pQacR<sup>1</sup> | 2.0 | 1.6 | 0.02 |

## Supplemental Information

**Supplemental Figures 1 - 5**

**Supplemental Tables 1 - 6**

**Supplemental Videos 1 - 10**

**Supplemental Model Information**

**References**

**Supplemental Figure 1.**
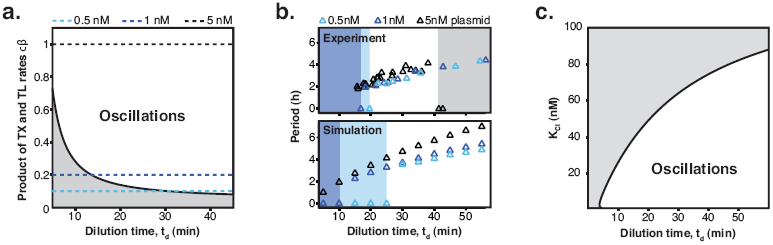
Oscillation parameter regime for a 3-node repressilator network in terms of dilution time. **a**, Transcription (TX) and translation (TL) rates supporting oscillations at different dilution times for a 3-node repressilator network. **b**, We experimentally studied the effect of varying transcription rates on the WT repressilator by measuring the range of dilution times that supported sustained oscillations. Transcription rates could be rapidly adjusted by varying DNA template concentrations of the repressilator plasmid. For different DNA template concentrations, oscillations occurred in different ranges of dilution times. Markers at a period of 0 h indicate a stable steady state, and shaded regions highlight dilution times that did not support oscillations for a specific DNA template concentration. A simulation of the repressilator network produced similar results but did not capture loading effects on the biosynthetic machinery for high DNA template concentrations. **c**, Increasing the K_M_ value of one repressor, as for CI repressor in the O_R_2* repressilator version, reduces the range of dilution rates that support oscillations as indicated by our experimental results (Fig. 2c) (see Supplementary Information for details on model).

**Supplemental Figure 2.**
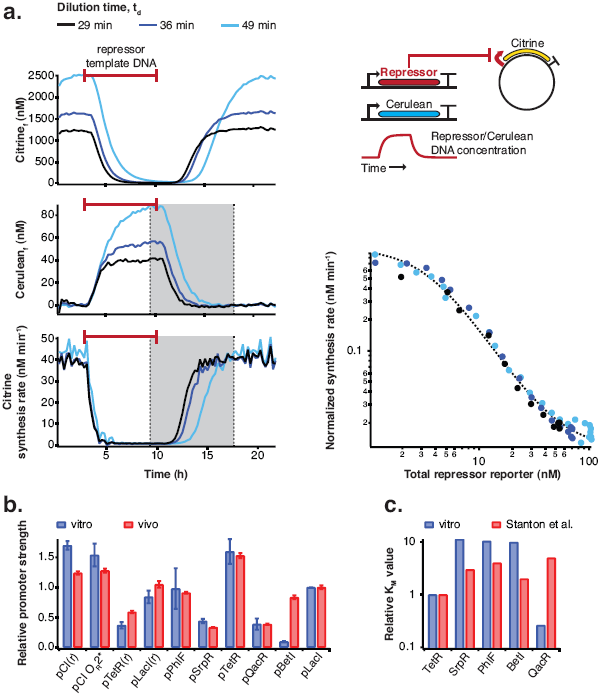
Repressor characterization. **a**, Transfer functions of the repressor – promoter pairs were determined using the cell-free framework as described in the Methods. Shown are experimental results and analysis using LacI - pLacI(r) as an example. Synthesis rates from the promoter of interest could be followed by Citrine fluorescence. Varying repressor template DNA concentration over time allowed us to determine synthesis rates at different repressor concentrations. Cerulean was co-expressed with the repressor and served as reporter for repressor concentration. Transfer functions were obtained by plotting Citrine synthesis rates from highest to lowest repressor concentration (grey shaded area) against total Cerulean concentration and were identical for different dilution times set in the nano-reactor device. **b**, Comparison of relative promoter strengths (v_max_), determined *in vitro* and *in vivo*. pCI(r), pTetR(r), and pLacI(r) are from^2^; pTetR is from^1^ and pLacI from^3^. Error bars indicate standard deviations of three replicates. **c**, Comparison of KM values measured *in vitro* in this study with KM values determined *in vivo* by Stanton et al.^1^. K_M_ values were normalized to the K_M_ of TetR.

**Supplemental Figure 3.**
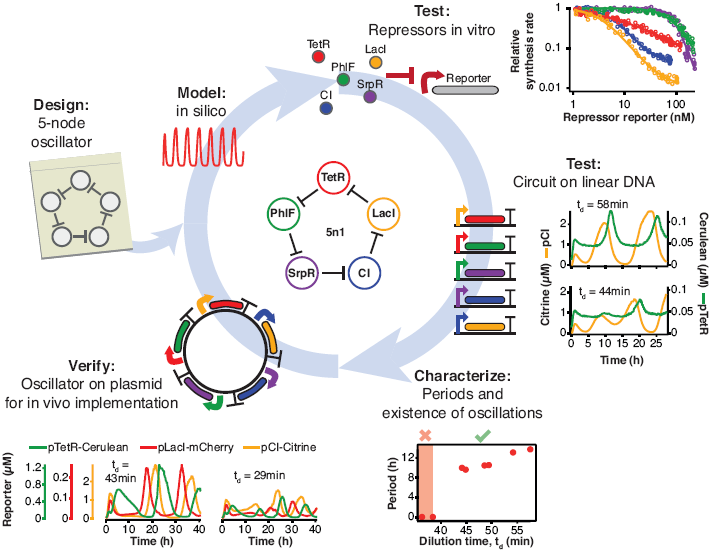
Engineering a 5-node negative feedback oscillator using the cell-free framework. A novel network architecture, which shows the intended behavior *in silico* is first assembled on linear DNA using *in vitro* characterized parts. Initial circuit testing on linear DNA is advantageous because: i) linear DNA can be synthesized in a few hours, ii) it allows rapid testing of multiple circuit variants, iii) and allows expression strengths of network components to be easily tuned by varying their relative concentrations. A functional circuit can then be further characterized to identify parameter ranges that support the desired behavior and to experimentally test hypotheses. If an *in vivo* implementation is intended, the cloned plasmids are verified for correct function *in vitro* before *in vivo* implementation.

**Supplemental Figure 4.**
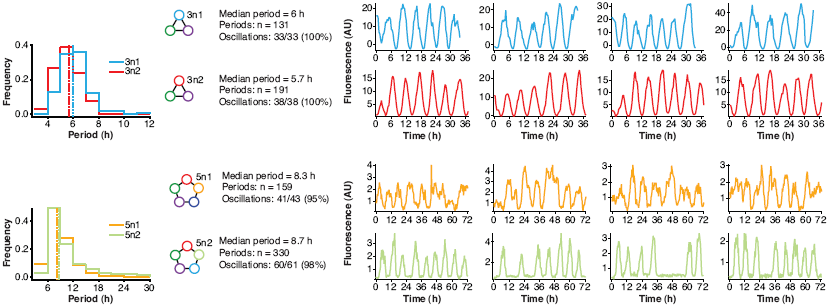
Robust oscillations of 3-node and 5-node oscillators *in vivo*. 3-node (top) and 5-node networks (bottom) oscillate with periods that depend on the network size *in vivo*. Shown are the distributions of observed period lengths with medians indicated by dashed lines. Both 3-node and 5-node networks exhibited robust oscillation with all growing cells oscillating for the 3-node networks and more than 95% of growing cells oscillating for the 5-node networks (defined as at least two distinct oscillation peaks per trace). Shown are four example traces for all oscillators in addition to the ones shown in Fig. 4a-b. Both 3-node networks were analyzed using a strong pPhlF sfGFP-ssrA reporter and the two 5-node networks were analyzed using a weak pPhlF sfGFP-ssrA reporter.

**Supplemental Figure 5.**
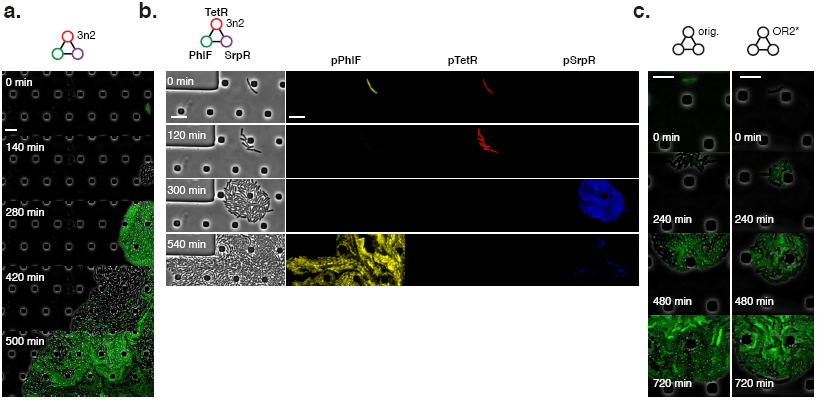
Population level synchronization of 3-node oscillators *in vivo*. **a**, 3n2 oscillator displays phase synchrony *in vivo*. 3n2 is run under a strong pPhlF sfGFP-ssrA reporter in the CellASIC microfluidic device. b, 3n2 displays phase synchrony observing 3 reporters simultaneously. Reporters are a strong pPhlF Citrine-ssrA, pTetR mCherry-ssrA, and pSrpR Cerulean-ssrA. Shown is one oscillation cycle. **c**, Original repressilator and O_R_2* repressilator do not show phase synchrony. These are run under pTetR(r)-eGFP(ASV) in M9 minimal media; oscillations were not supported in LB. All scale bars: 10 μm.

**Supplemental Table 1.** Strains used in this study.

| Name | *E. coli* type | Resistance | Notes |
| --- | --- | --- | --- |
| Rosetta2 | JS006 | cmR |  |
| pZS1 + pZE21-GFP(AAV) | JS006 | kanR ampR |  |
| pET21a(+)-Histag-Citrine | BL21-DE3 | ampR |  |
| pET21a(+)-Histag-Cerulean | BL21-DE3 | ampR |  |
| pET21a(+)-Histag-mCherry | BL21-DE3 | ampR |  |
| 3n1 + pPhIF-BCD20-sfGFP-ssrA(LAA) | JS006 | kanR ampR | minimize passages |
| 3n1 + pPhIF-BCD22-sfGFP-ssrA(LAA) | JS006 | kanR ampR | minimize passages |
| 3n2 + pPhIF-BCD20-sfGFP-ssrA(LAA) | JS006 | kanR ampR | minimize passages |
| 3n2 + pPhIF-BCD22-sfGFP-ssrA(LAA) | JS006 | kanR ampR | minimize passages |
| 5n1 + pPhIF-BCD20-sfGFP-ssrA(LAA) | JS006 | kanR ampR | cells unhealthy, minimize passages |
| 5n1 + pPhIF-BCD22-sfGFP-ssrA(LAA) | JS006 | kanR ampR | minimize passages |
| 5n2 + pPhIF-BCD20-sfGFP-ssrA(LAA) | JS006 | kanR ampR | minimize passages |
| 5n2 + pPhIF-BCD22-sfGFP-ssrA(LAA) | JS006 | kanR ampR | minimize passages |
| 3n2 + 3-color BCD20 reporter, pPhIF/pSrpR/pJ23119-tetO | JS006 | kanR cmR | minimize passages |
| pZS1 + pZE21-eGFP(ASV) | JS006 | kanR ampR |  |
| pZS1 w/ OR2* mutation + pZS21-eGFP(ASV) | JS006 | kanR ampR |  |

**Supplemental Table 2.**
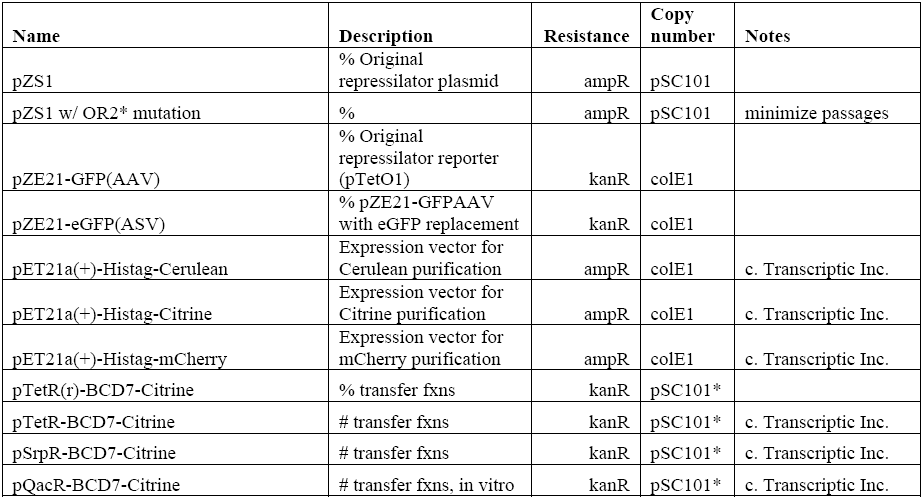
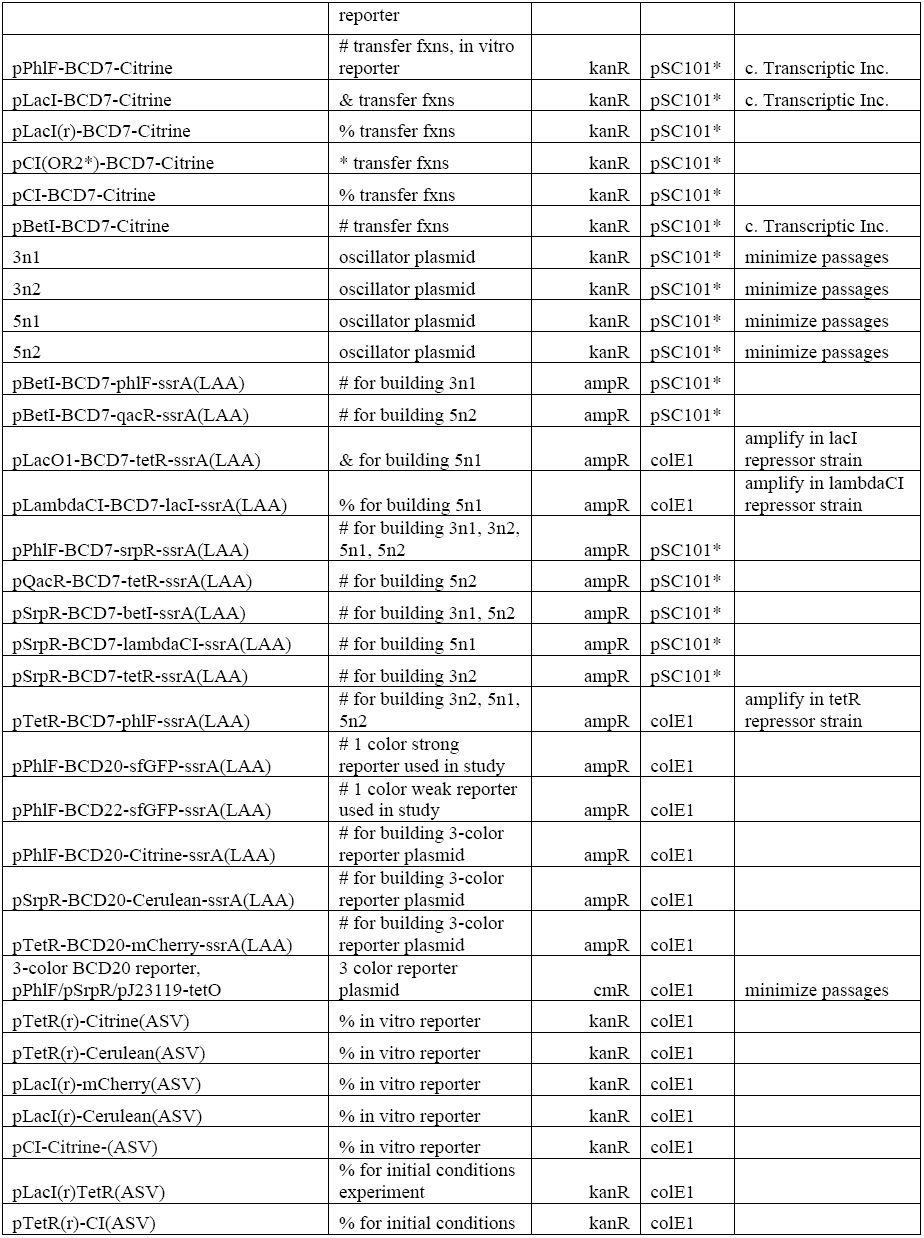
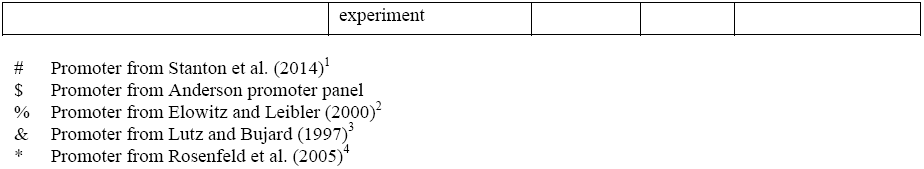
Plasmids used in this study.

**Supplemental Table 3.**
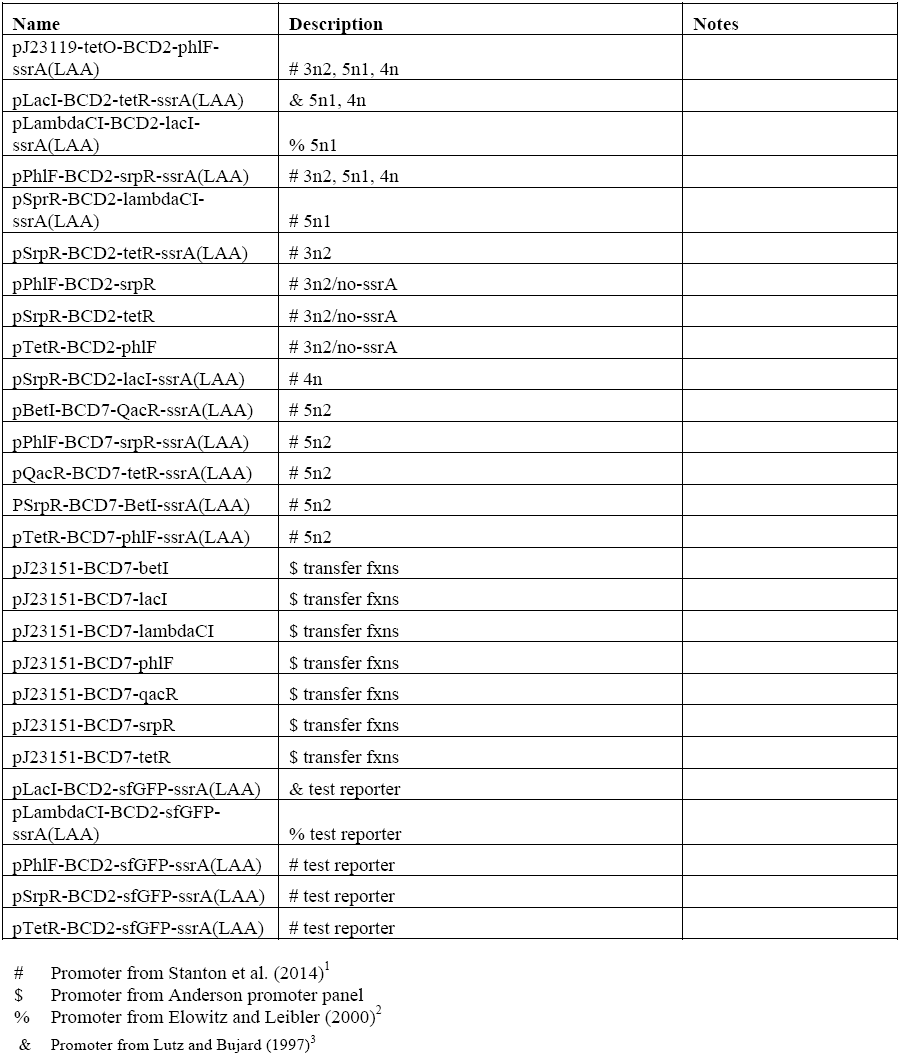
Linear DNAs used in this study.

| Name | Description | Notes |
| --- | --- | --- |
| pJ23119-tetO-BCD2-phlF-ssrA(LAA) | # 3n2, 5n1, 4n |  |
| pLacI-BCD2-tetR-ssrA(LAA) | & 5n1, 4n |  |
| pLambdaCI-BCD2-lacI-ssrA(LAA) | % 5n1 |  |
| pPhlF-BCD2-srpR-ssrA(LAA) | # 3n2, 5n1, 4n |  |
| pSprR-BCD2-lambdaCI-ssrA(LAA) | # 5n1 |  |
| pSrpR-BCD2-tetR-ssrA(LAA) | # 3n2 |  |
| pPhlF-BCD2-srpR | # 3n2/no-ssrA |  |
| pSrpR-BCD2-tetR | # 3n2/no-ssrA |  |
| pTetR-BCD2-phlF | # 3n2/no-ssrA |  |
| pSrpR-BCD2-lacI-ssrA(LAA) | # 4n |  |
| pBetI-BCD7-QacR-ssrA(LAA) | # 5n2 |  |
| pPhlF-BCD7-srpR-ssrA(LAA) | # 5n2 |  |
| pQacR-BCD7-tetR-ssrA(LAA) | # 5n2 |  |
| PSrpR-BCD7-BetI-ssrA(LAA) | # 5n2 |  |
| pTetR-BCD7-phlF-ssrA(LAA) | # 5n2 |  |
| pJ23151-BCD7-betI | \$ transfer fxns | |
| pJ23151-BCD7-lacI | \$ transfer fxns | |
| pJ23151-BCD7-lambdaCI | \$ transfer fxns | |
| pJ23151-BCD7-phlF | \$ transfer fxns | |
| pJ23151-BCD7-qacR | \$ transfer fxns | |
| pJ23151-BCD7-srpR | \$ transfer fxns | |
| pJ23151-BCD7-tetR | \$ transfer fxns | |
| pLacI-BCD2-sfGFP-ssrA(LAA) | & test reporter |  |
| pLambdaCI-BCD2-sfGFP-ssrA(LAA) | % test reporter |  |
| pPhlF-BCD2-sfGFP-ssrA(LAA) | # test reporter |  |
| pSrpR-BCD2-sfGFP-ssrA(LAA) | # test reporter |  |
| pTetR-BCD2-sfGFP-ssrA(LAA) | # test reporter |  |

- # Promoter from Stanton et al. (2014)<sup>1</sup> \$ Promoter from Anderson promoter panel % Promoter from Elowitz and Leibler (2000)<sup>2</sup>
& Promoter from Lutz and Bujard (1997)<sup>3</sup>

**Supplemental Table 4.** DNA concentrations used in experiments.

| Experiment | DNA and concentration | Type of DNA |
| --- | --- | --- |
| **Repressilator orig./O<sub>R</sub>2\*, 3color** | Repressilator pZS1 or pZS1 w/ O<sub>R</sub>2* mutation, 0.5 nM (if not otherwise indicated) | Plasmid |
|  | pTetR(r)-Cerulean(ASV), 5 nM | Plasmid |
|  | pLacI(r)-mCherry(ASV), 5 nM | Plasmid |
|  | pCI-Citrine(ASV), 5 nM | Plasmid |
| **Repressilator, initial conditions** | **Reaction in nano-reactor:** |  |
|  | Repressilator pZS1, 5 nM | Plasmid |
|  | pLacI(r)-Cerulean(ASV), 5 nM | Plasmid |
|  | pTetR(r)-Citrine-(ASV), 5 nM | Plasmid |
|  | **Pre-synthesis reaction (CI):** |  |
|  | pTetR(r)-Citrine(ASV), 5 nM | Plasmid |
|  | pTetR(r)-CI(ASV), 5 nM | Plasmid |
|  | **Pre-synthesis reaction (TetR):** |  |
|  | pLacI(r)-Cerulean(ASV), 5 nM | Plasmid |
|  | pLacI(r)TetR(ASV), 5 nM | Plasmid |
| **Response curve measurements** | Promoter plasmid: pXXX-BCD7-Citrine, 1 nM | Plasmid |
|  | Repressor template: pJ23151-BCD7-XXX, 0-2.5 nM | Linear |
|  | Repressor reporter: pJ23151-BCD7-Cerulean, 0-2.5 nM | Linear |
| **3n1** | 3n1 oscillator plasmid, 5 nM | Plasmid |
|  | pPhlF-BCD7-Citrine, 2.5 nM | Plasmid |
| **3n2** | pJ23119-tetO-BCD2-phlF-ssrA(LAA), 1.5 nM | Linear |
|  | pPhlF-BCD2-srpR-ssrA(LAA), 12 nM | Linear |
|  | pSrpR-BCD2-tetR-ssrA(LAA), 24 nM | Linear |
|  | pTetR(r)-Cerulean(ASV), 5 nM | Plasmid |
| **3n2/no-ssrA** | pJ23119-tetO-BCD2-phlF, 1.5 nM | Linear |
|  | pPhlF-BCD2-srpR, 12 nM | Linear |
|  | pSrpR-BCD2-tetR, 24 nM | Linear |
|  | pTetR(r)-Cerulean(ASV), 5 nM | Plasmid |
| **4n** | pJ23119-tetO-BCD2-phlF-ssrA(LAA), 0.75 nM | Linear |
|  | pLacI-BCD2-tetR-ssrA(LAA), 6 nM | Linear |
|  | pPhlF-BCD2-srpR-ssrA(LAA), 6 nM | Linear |
|  | pSrpR-BCD2-lacI-ssrA(LAA), 12 nM | Linear |
|  | pTetR(r)-Cerulean(ASV), 2.5 nM | Plasmid |
|  | pLacI(r)-mCherry(ASV), 2.5 nM | Plasmid |
|  | pPhlF-BCD7-Citrine, 2.5 nM | Plasmid |
| **5n1** | pJ23119-tetO-BCD2-phlF-ssrA(LAA), 1.1 nM | Linear |
|  | pLacI-BCD2-tetR-ssrA(LAA), 16.8 nM | Linear |
|  | pLambdaCI-BCD2-lacI-ssrA(LAA), 1.4 nM | Linear |
|  | pPhlF-BCD2-srpR-ssrA(LAA), 5.6 nM | Linear |
|  | pSprR-BCD2-lambdaCI-ssrA(LAA), 11.2 nM | Linear |
|  | pCI-Citrine(ASV), 3 nM | Plasmid |
|  | pTetR(r)-Cerulean(ASV), 2.5 nM | Plasmid |
| **5n1, plasmid DNA** | 5n1 oscillator plasmid, 5 nM | Plasmid |
|  | pTetR(r)-Cerulean(ASV), 5 nM | Plasmid |
|  | pLacI(r)-mCherry(ASV), 5 nM | Plasmid |
|  | pCI-Citrine(ASV), 5 nM | Plasmid |
| **5n2** | pBetI-BCD7-QacR-ssrA(LAA), 1 nM | Linear |
|  | pPhlF-BCD7-srpR-ssrA(LAA), 12 nM | Linear |
|  | pQacR-BCD7-tetR-ssrA(LAA), 4 nM | Linear |
|  | pSrpR-BCD7-BetI-ssrA(LAA), 24 nM | Linear |
|  | pTetR-BCD7-phlF-ssrA(LAA), 4 nM | Linear |
|  | pTetR(r)-Cerulean(ASV), 2.5 nM | Plasmid |
|  | pQacR-BCD7-Citrine, 2.5 nM | Plasmid |

### Supplemental Video 1

3-color *in vitro* run of repressilator at t_d_ = 47 min. Shown on the right are individual channels of pCI-Citrine-ssrA, pTetR-Cerulean-ssrA, and pLacI-mCherry-ssrA. These are combined in the composite at the left.

### Supplemental Video 2

3n1 in mother machine, single trap. 3n1 using a pPhlF-BCD20-sfGFP-ssrA (strong) reporter is run in the mother machine at 29°C in LB.

### Supplemental Video 3

3n2 in mother machine, single trap. 3n2 using a pPhlF-BCD20-sfGFP-ssrA (strong) reporter is run in the mother machine at 29°C in LB.

### Supplemental Video 4

Run of a panel of 3n2 oscillators in mother machine, using pPhlF-BCD20-sfGFP-ssrA (strong) reporter.

### Supplemental Video 5

5n1 in mother machine. 5n1 using a pPhlF-BCD22-sfGFP-ssrA (weak) reporter is run in the mother machine at 29°C in LB.

### Supplemental Video 6

5n2 in mother machine. 5n2 using a pPhlF-BCD22-sfGFP-ssrA (weak) reporter is run in the mother machine at 29°C in LB.

### Supplemental Video 7

Run of a panel of 5n2 oscillators in mother machine, using pPhlF-BCD22-sfGFP-ssrA (weak) reporter.

### Supplemental Video 8

3n1 in CellASIC. 3n1 using a pPhlF-BCD20-sfGFP-ssrA (strong) reporter is run in CellASIC. This video corresponds to Fig. 4c. Conditions: 29°C, LB media, 12.5% lamp intensity, 200 ms exposure, 2 psi flow rate.

### Supplemental Video 9

3n2 in CellASIC. 3n2 using a pPhlF-BCD20-sfGFP-ssrA (strong) reporter is run in CellASIC. This video corresponds to Supp. Fig. 5a. Conditions: 29°C, LB media, 12.5% lamp intensity, 200 ms exposure, 2 psi flow rate.

### Supplemental Video 10

3n2 with 3-color output in mother machine. 3n2 using a pPhlF-BCD20-Citrine-ssrA, pTetR-BCD20-mCherry-ssrA, pSrpR-Cerulean-ssrA (strong) reporter is run in CellASIC. This video corresponds to Supp. Fig. 5b. Conditions: 29 C, LB media, 12.5% lamp intensity, 200 ms exposure for Citrine and Cerulean (500 ms for mCherry), 2 psi flow rate.

## Supplemental Model Information

### Mathematical model

We consider an *n*-node negative cyclic feedback biocircuit and denote the genes, mRNAs and proteins by *G*_1_, *G*_2_, …, *G*_*n*_, *M*_1_, *M*_2_, …, *M*_*n*_ and *P*_1_*P*_2_, …, *P*_*n*_, respectively. Let *r*_*i*_(*t*) and *p*_*i*_(*t*) denote the concentrations of mRNA *M*_*i*_ and protein *P*_*i*_, respectively. For example, the novel 3-node ring oscillator in Fig. 3b is defined by *n* = 3, *r*_1_ (*t*) = [Betl mRNA], *r*_2_(*t*) = [PhlF mRNA], *r*_3_(*t*) *=* [SrpR mRNA], *p*_1_(*t*) = [Betl protein], *p*_2_ (*t*) = [PhlF protein] and *p*_3_(*t*) = [SrpR protein].

Our mathematical model considers transcription, translation and degradation of mRNA and protein molecules as summarized in Supp. Table 5, where *a*_*i*_ and *b*_*i*_ represent the degradation rates of *M*_*i*_ and *P*_*i*_, respectively, and *c*_*i*_ and *β*_*i*_ are the translation and transcription rates. The constants *K*_*i*−1_ and *v*_*i*_ are the Michaelis-Menten constant and the Hill coefficient associated with the protein *P*_*i*−1_ and the corresponding promoter on gene *G*_*i*_. We hereafter use subscripts 0 and *n* + 1 as the substitutes of *n* and 1, respectively, to avoid notational clutter.

Using the law of mass action and the quasi-steady state approximation, the dynamics of the mRNA and protein concentrations can be modeled by the following ordinary differential equations (ODE).

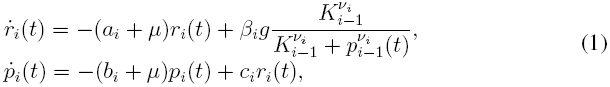

where *i* =1, 2, …, *n*, and *g* is the concentration of the circuit plasmid. The constant *μ* is the dilution rate of mRNA and proteins by the microfluidic device. The dilution time of the microfluidic device is defined by

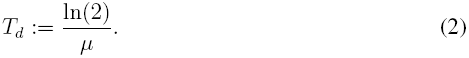

### Numerical prediction of the oscillation period

The ODE model (1) was numerically simulated using `ode45` solver of MATLAB R2013b to obtain qualitative insight into the period as well as the oscillatory parameter regime (Fig. 3f and Supp. Fig. 1b). The parameters in Supp. Table 6 were used for the simulations.

**Supplemental Table 5.** Stoichiometry and reaction rates.

| Description | Reaction | Reaction rate |
| --- | --- | --- |
| Transcription of $M_i$ | $G_i + P_{i-1} \rightarrow G_i + P_{i-1} + M_i$ | $\beta_i \frac{K_{i-1}^{\nu_i}}{K_{i-1}^{\nu_i} + p_{i-1}^{\nu_i}}$ |
| Translation of $M_i$ | $M_i \rightarrow M_i + P_i$ | $c_i r_i$ |
| Degradation of $M_i$ | $M_i \rightarrow \emptyset$ | $a_i r_i$ |
| Degradation of $P_i$ | $P_i \rightarrow \emptyset$ | $b_i p_i$ |

**Supplemental Table 6.**
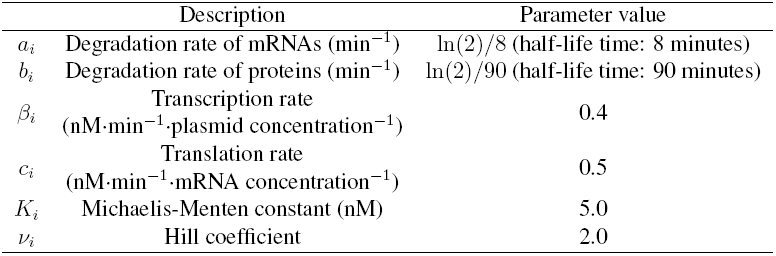
Parameters used for simulations.

|  | Description | Parameter value |
| --- | --- | --- |
| $a_i$ | Degradation rate of mRNAs ($\text{min}^{-1}$) | $\ln(2)/8$ (half-life time: 8 minutes) |
| $b_i$ | Degradation rate of proteins ($\text{min}^{-1}$) | $\ln(2)/90$ (half-life time: 90 minutes) |
| $\beta_i$ | Transcription rate<br>($\text{nM} \cdot \text{min}^{-1} \cdot \text{plasmid concentration}^{-1}$) | 0.4 |
| $c_i$ | Translation rate<br>($\text{nM} \cdot \text{min}^{-1} \cdot \text{mRNA concentration}^{-1}$) | 0.5 |
| $K_i$ | Michaelis-Menten constant (nM) | 5.0 |
| $\nu_i$ | Hill coefficient | 2.0 |

The plasmid concentration *g* was set as *g* = 5.0 nM for Fig. 3f. The initial concentrations for the simulations were *r*_1_(0) = 30, *p*_1_(0) = 0 and *r*_*i*_(0) = *p*_*i*_(0) *=* 0 for *i* = 2, 3, …, *n*.

The period of oscillations was calculated based on the autocorrelation of the simulated protein concentration *p*_1_(*t*). More specifically, let

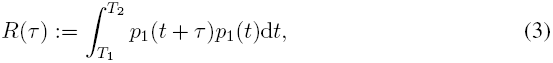

where *T*_1_ is a positive constant such that *p*_1_(*t*) is steady state at *t* = *T*_1_, and *T*_2_ is a sufficiently large constant compared to the period of oscillations. The period of oscillations *T*_period_ was determined by *T*_period_ = min_*τ*>0_ argmax_*τ*_*R*(*τ*). The simulation result is also consistent with the analytic estimation of the oscillation period in Hori *et al.^5^* in that the period increases monotonically with the dilution time *T*_*d*_.

### Parameter regions for oscillations

The parameter region for oscillations (Supp. Fig. 1a) was obtained based on the analysis result (Theorem 3) by Hori *et al.^5^*. Since parameter values do not depend on the subscript *i* as shown in Supp. Table 6, we remove the subscript *i* and define *a* := *a*_1_(= *a*_2_ = … *a*_*n*_). In the same way, we define *b*, *c*, *β*, *K* and *ν*.

It was shown that the protein concentrations *p*_*i*_ (*i* = 1, 2, …, *n*) oscillate if both of the following inequalities are satisfied^5^.

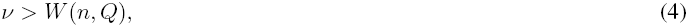

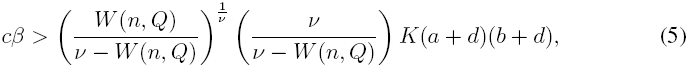

Where

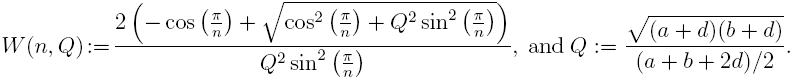

To obtain the parameter region in Supp. Fig. 1a, we substituted *n* = 3 and the parameters shown in Supp. Table 6 into the right-hand side of the inequality condition (5), then we varied *T*_*d*_(= ln(2)/*μ*) between 5 to 45. The inequality (4) was always satisfied for these parameters.

The parameter region of Supp. Fig. 1c was obtained by the local stability analysis of the model (1). The previous theoretical result^6^ showed that the model (1) has a unique equilibrium point and the protein concentrations *p*_*i*_ (*i* = 1, 2, …, *n*) show stable oscillations if the Jacobian matrix evaluated at the equilibrium point has an eigenvalue in the open right-half complex plane. Based on this result, we computed the Jacobian eigen-values with varying *K*_3_, which we denote by *K*_cI_, and *T*_*d*_. The values in Supp. Table 6 were used for the other parameters. The plasmid concentration was set as *g* = 5.0 nM in the computation.

